# Prolonged culture of primary human keratinocytes isolated from suction blisters

**DOI:** 10.1101/333831

**Authors:** Erik D. Anderson, Inka Sastalla, Noah J. Earland, Minai Mahnaz, Ian N. Moore, Francisco Otaizo-Carrasquero, Timothy G. Myers, Christopher A. Myles, Sandip K. Datta, Ian A. Myles

## Abstract

Keratinocytes are the most abundant cell type in the epidermis. They prevent desiccation and provide immunological and barrier defense against potential pathogens such as *Staphylococcus aureus* and *Candida albicans*. The study of this first line of immune defense is hindered by invasive isolation methods and insufficient techniques for long-term passage of primary keratinocytes *in vitro.* Primary keratinocytes have been successfully isolated from blister roofs induced by negative pressure, which separates the epidermis from the dermis *in vivo* in human subjects. This method allows collection of pure epidermal cells without dermal contamination in a minimally invasive manner. However, the isolated keratinocytes differentiate and senesce when cultured *in vitro* beyond five passages. Here, we present evidence that the Rho kinase (ROCK) inhibitor Y-27632 can be used to effectively increase the proliferative capabilities of keratinocytes isolated using the suction blister method, similar to what has been previously reported for primary keratinocytes isolated using alternative methods. We show that the increase in passage number is directly correlated to delayed differentiation, and that cells passaged long term with the inhibitor retain their ability to stratify in organotypic raft cultures and respond to cytokine treatment; additionally, the late passage cells have a heterogeneous mix of differentiated and non-differentiated cells which may be predicted by a ratio of select differentiation markers. The described method presents a minimally invasive procedure for keratinocyte isolation and prolonged culture that allows analysis of keratinocyte function in both healthy volunteers and patients with dermatologic diseases.

## Introduction

The skin consists of three layers: the epidermis, dermis, and subcutaneous tissue. Keratinocytes are the most abundant cell type in the epidermis and, among other functions, prevent desiccation while providing barrier defense against bacterial, viral, and fungal pathogens such as *Staphylococcus aureus* [1] and *Candida albicans* [2]. Research into these important functions of primary keratinocytes is hindered by invasive isolation methods and limited availability of techniques for long-term *ex vivo* culture.

The acquisition of primary keratinocytes from the epidermal roof of suction blisters overcomes some limitations of standard keratinocyte isolation methods. Collection of suction blister keratinocytes (SBK) is accomplished by utilizing negative pressure to separate the epidermis from the dermis *in vivo*. The effect is the formation of a blister composed of epidermal cells that can be excised and treated with trypsin to dissociate the cells [3]. This isolation method holds advantages over others — such as keratinocyte isolation from skin biopsies, neonatal foreskins, and whole skin explants from surgical procedures — because it is minimally invasive and results in the isolation of pure epidermal cells without any dermal contamination [4]. The technique also facilitates the study of keratinocytes from patients diagnosed with diseases after infancy, and importantly, it is not confined to a specific gender. However, SBK are difficult to propagate for an extended period of time due to the propensity of the cells to terminally differentiate and senesce in culture [5].

The Rho family of small GTP-binding proteins affect a wide range of cell functions, such as assembly of the actin cytoskeleton, cell proliferation, and motility [6]. Two Rho associated protein kinases (ROCKs) in the mammalian system are downstream effectors of Rho, and their activation is correlated with stress-fiber formation and cellular contraction [7]. The chemical Y-27632 inhibits both ROCK isoforms [8] and prevents dissociation-induced apoptosis in human embryonic stem cells in culture [9]. Chapman *et al.* showed that culture in the presence of Y-27632 effectively immortalizes human neonatal foreskin keratinocytes (HFK) and adult vaginal and ectocervical keratinocytes isolated from hysterectomy tissue [10]. They reported that the keratinocytes passaged with Y-27632 retain normal karyotypes and morphologies, as well as the ability to stratify in organotypic raft cultures after removal of the inhibitor.

In this study, we show that *in vitro* cultivation of SBK in the presence of the ROCK inhibitor Y-27632 increases their proliferative capacity and delays terminal differentiation, consistent with what has been reported for keratinocytes derived from other sources [10, 11]. Compared to these previous reports, we found a higher presence of differentiated keratinocytes, including corneocytes (terminally differentiated keratinocytes) in cultures of cells passaged more than 40 days with Y-27632. However, these SBK passaged long term with the inhibitor retained their ability to stratify in organotypic raft culture and to respond to interleukin (IL)-17 and interferon (IFN)-γ. Additionally, retention of these functional properties inversely correlated with the differentiation state of the culture as measured by the ratio of involucrin (IVL) to keratin (KRD. T)-14 protein expression. These studies identify propagation of SBK in the presence of Y-27632 as a viable and valuable method to investigate the function of primary human keratinocytes.

## Results

### Y-27632 increases the proliferative capacity of SBK in culture

Chapman *et al.* reported previously that keratinocytes isolated from neonoatal foreskin and surgical explants can be indefinitely cultured in the presence of the ROCK inhibitor Y-27632, achieving 150 passages over a period of 500 days [10]. We tested whether similar results could be achieved with keratinocytes isolated using the suction-blister method.

Many suction blister studies employ mono-chamber devices, often utilizing syringes with the plungers removed and attached to vacuum pumps [12, 13]. These chambers can be cumbersome, cannot assure equal pressure between independent chambers, and/or lack the ability to monitor the pressure during the application. Other studies have deployed multi-chamber devices but with the chambers aligned in series [4, 14]. Unfortunately, these devices produce heterologous blisters due to the barotrauma gradient between the wells closest to the vacuum source and those farthest from it. Therefore, we developed a 3D printed chamber that could evenly distribute the barometric pressure across eight independent, 8mm blisters (S1 Appendix) and attaches to commercially available, micro-derm abrasion devices with titratable vacuum pressure. This approach assured reproducible barometric pressure between different subjects and each well on a given participant. Furthermore, the total yield of skin was 401.6 mm^2^ (8 × π4^2^), the equivalent of a 22.6 mm biopsy.

Keratinocytes were isolated from healthy volunteers using our device (Fig 1A) and passaged in the presence or absence of Y-27632. The inhibitor increased the terminal population doubling of SBK from 1.8 after 22 days to 14.7 after 70 days (Fig 1B), and generated approximately 6X as many cells in culture (Fig 1C). Thus, addition of Y-27632 can be used to increase cell yield and proliferation of SBK, but the culture could not be extended indefinitely as indicated by the plateu of the nonlinear curve. In contrast, HFK exhibited greater population doubling in the presence or absence of Y-27632 (Fig 1B, S1 Fig), as previously reported [10, 11].

**Fig 1.**
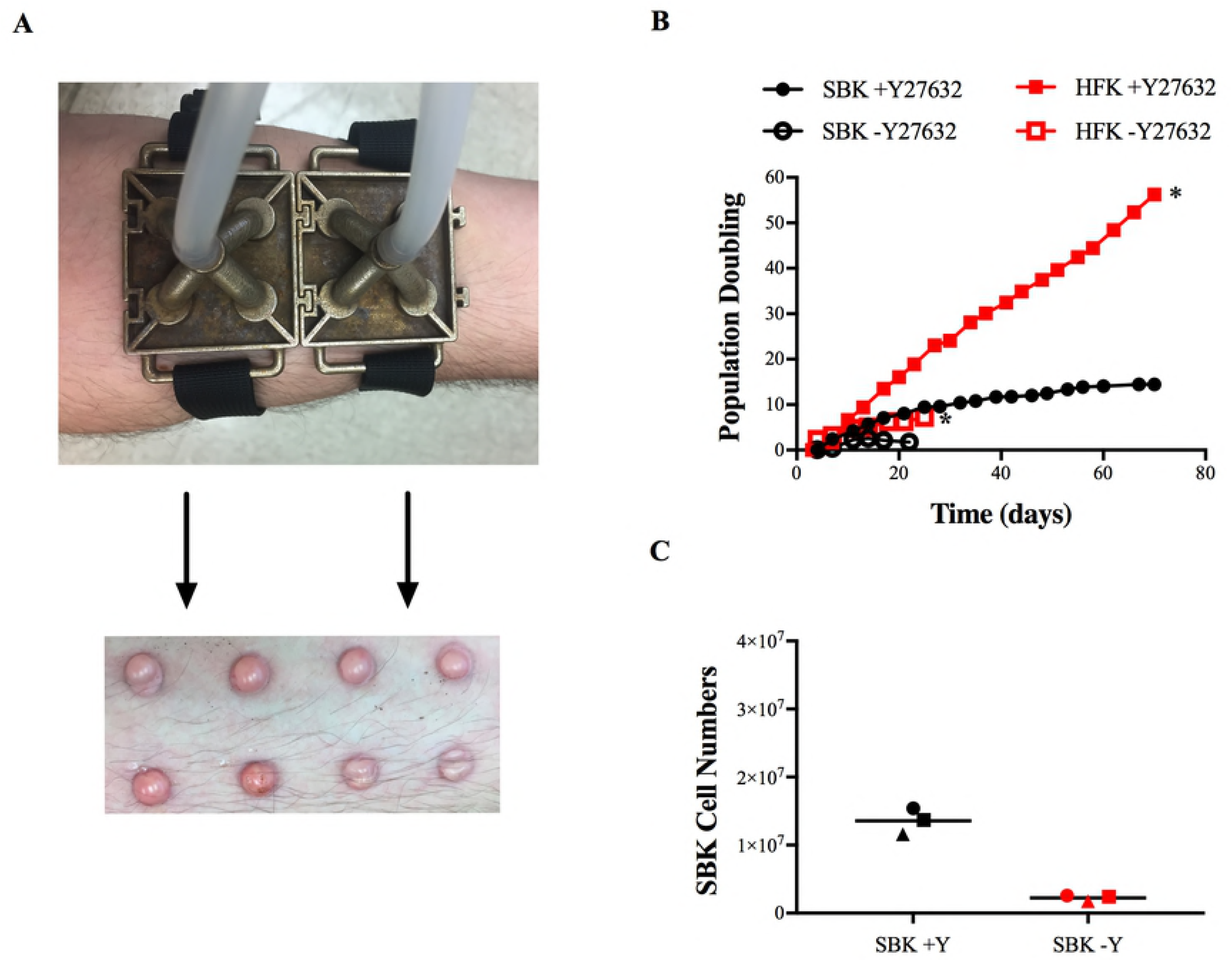
Y-27632 increases the proliferation of SBK in culture. (A) Application of the suction-blister device to the forearm (left), and the appearance of epidermal blisters after 2 hours of suction (right). (B) The population doubling of SBK (black, n=3 patient lines, average and standard deviation of the cultures are shown) and HFK (red, n=1) passaged with (+Y) or without (-Y) the ROCK inhibitor Y-27632. The asterix indicates when HFK cultures were intentionally stopped. SBK cultures were passaged to senescence. (C) Total SBK cell numbers generated from the +Y (black) and -Y (red) conditions after 70 and 22 days respectively (each shape represents a distinct source subject (n=3), bar indicates mean.)

### Y-27632 delays terminal differentiation of SBK

To characterize the effect of Y-27632 on SBK, we isolated RNA from early pass cells cultured either with or without the inhibitor and analyzed gene expression using RNAseq. Y-27632 has been previously described to inhibit differentiation of adult keratinocytes derived from skin biopsies, and microarray analysis found some of the genes within the epidermal differentiation complex (EDC) to be downregulated [11]. The EDC is a group of genes located on chromosome 1q21 that encode proteins important for keratinocyte differentiation into corneocytes [15]. We identified over 1,100 differentially expressed genes with ±2.6 log_2_ fold change in the group containing Y-27632 (+Y) as compared to the group lacking it (-Y) (S2 Appendix), and of these genes 929 were downregulated (Fig 2A). Of the downregulated genes, approximately 20% of the epidermal differentiation complex genes were identified, with more than half previously reported to be downregulated in keratinocytes derived from adult skin biopsies (Fig 2B).

**Fig 2.**
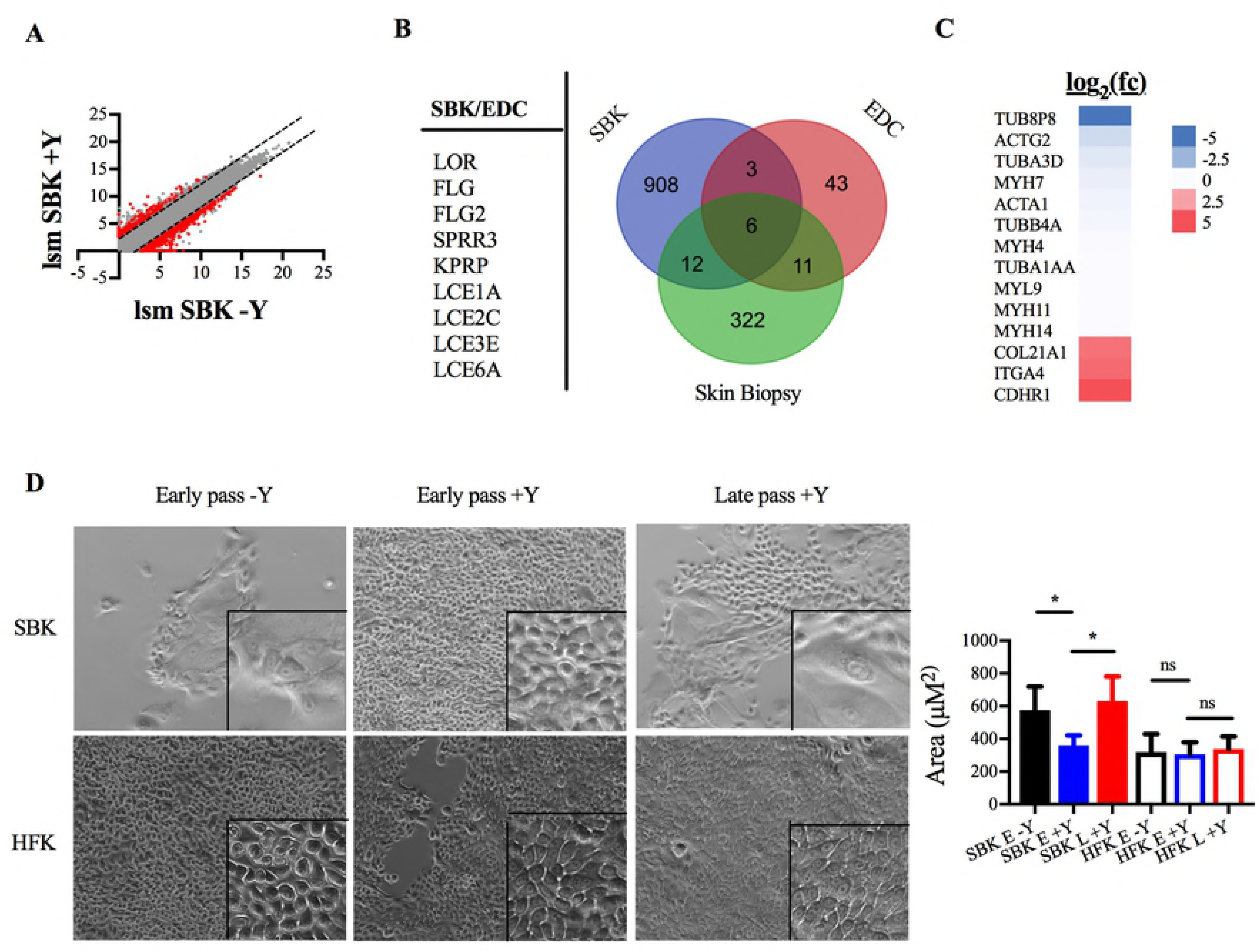
Y-27632 delays terminal differentiation in SBK. RNA-seq analysis was performed on RNA isolated from SBK grown in the absence (-Y; passages 4, 5, and 6) and presence (+Y; passages 5 and 6) of Y-27632. (A) A plot of the log_2_ least square mean of all the genes from each culture condition. The dotted lines indicate genes with greater than 2.6 fold change; the red dots indicate the subset of those with an adjusted p value < 0.5. (B) The Venn diagram compares the overlap between significantly downregulated genes previously described in adult skin biopsy keratinocytes [11], SBK genes identified in this study, and genes associated with the epidermal differentiation complex (EDC). (C) Heatmap presenting the log_2_ fold change of identified cytoskeletal and adhesion protein genes. (D) Left, monolayers of early (passage 4; 12 days) or late (passage 14; 49 days) SBK and HFK after removal of the fibroblast feeder layer; scale bar = 100μM. Boxed areas represent 6X magnification. Right, area of cytoplasm in SBK (n=6/condition) or HFK (n=3/condition) measured from 25 cells for each image; ^*^, p <0.0001 (ANOVA with Sidak’s post test).

We also identified members of the tubulin, actin, and myosin gene families to be downregulated (Fig 2C), consistent with previous reports characterizing an inhibitory effect of Y-27632 on myosin-light chain phosphorylation [16] and stress fiber disassembly [17]. Additionally, genes involved in the attachment of basal keratinocytes to the basement membrane were upregulated (Fig 2C), a finding that may explain the increase in cell adherence and clumping observed for SBK incubated with Y-27632 immediately after blister isolation (S2 Figure, S1 Movie).

The eventual senescence of SBK despite the observed downregulation of differentiation-associated genes in the presence of Y-27632 prompted us to investigate whether any morphological indications of terminal differentiation could be detected in late passage keratinocytes. Based on proliferation kinetics (Fig 1B), we defined early passage cells as those harvested between passage 4-6, and late passage SBK as those harvested between passage 12-14. Increased cell size indicates keratinocyte differentiation towards the terminal anuclear corneocyte [18]. Thus, to assess for the presence of differentiated cells we measured the cytoplasmic area of SBK, and for comparison HFK, at early and late passages in the presence and absence of Y-27632 (Fig 2D). We observed that the average area of early passage SBK cultured without the inhibitor was significantly increased compared to those cultured with the inhibitor (Fig 2D). A significant difference was also detected between early and late passage SBK cultured with Y-27632. In contrast, HFK did not demostrate any significant morphologic changes with passage, consistent with the linear population doubling data in Fig 1B that suggest a lack of differentiation and continued proliferation even after 70 days in culture. Taken together, these data suggest that Y-27632 delays, but does not block, the terminal differentiation of SBK, in contrast to its immortalizing effect on HFK.

### Late passage SBK stratify into an epidermis

Next, we tested whether the increase in differentiating keratinocytes in the late passage cultures affected the functionality of the cells. One such functional measure is the ability of the cells to stratify into an epithelium because this process requires a dynamic and complex pattern of protein expression for proper differentiation and movement up the strata [19, 20]. For example, keratin-14 (KRT14) is an intermediate fillament protein expressed in keratinocytes that make up the stratum basale layer of the epidermis [21, 22], whereas Involucrin (IVL) is expressed as the cells differentiate to form the suprabasal layers [22, 23]. Early and late passage cells were grown as previously described [24] on collagen rafts and the stratification patterns were compared. H&E staining revealed less distinct layering in late passage SBK, with diminished appearance of the stratum corneum and resulted in a less prominent stratum granulosum layer (Fig 3A, B). Additionally, cornified cells were present in the subcorneal layer (Fig 3B, arrows); this finding was confirmed by immunofluorescent nuclear staining through the observed absence of nuclei in the lower strata layers of the late pass cells (Fig 3C, D). This indicates denuclearized cells, a hallmark of cornified terminally differentiated keratinocytes [25]. Despite these changes in the late passage cultures, both early and late passage cells retained normal stratification patterns for KRT14 (basal layers) and IVL (upper layers) (Fig 3C, D). However, while the localization of IVL and KRT14 was preserved in 3D cultures, the ratio of total IVL:KRT14 protein expression in the cultures increased with passage (S3 Fig). This further suggests SBK continue to undergo differentiation with each passage, and may indicate this ratio could be used as a measure of the proportion of differentiated cells within the culture.

**Fig 3.**
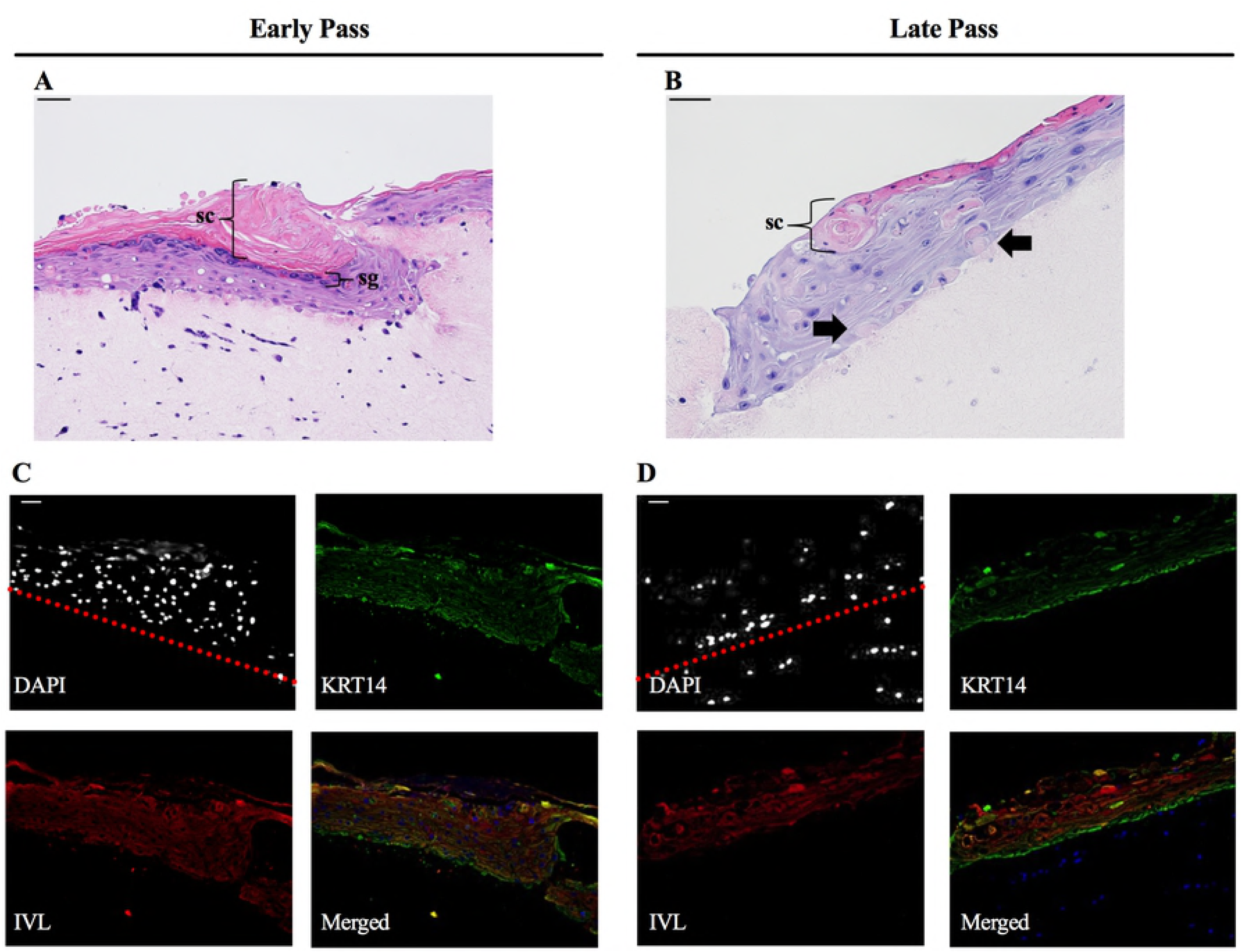
Late passage SBK stratify into an epidermis with increased numbers of cornified cells present in the subcorneal layers. Hematoxylin and eosin stain (A, B) and immunofluorescence stain (C, D; green, keratin 14; red, involucrin; blue, DAPI) of organotypic raft cultures derived from early and late passage keratinocytes. DAPI image presented in black and white to better distinguish the nuclei, and the red dotted line delineates where the epidermis begins. Nuclei below the line are from the 3T3 fibroblasts seeded in the collagen rafts. sc=stratum corneum, sg=stratum granulosum, scale bar = 100μM for both H&E and IF photos.

### Functional characterization of early and late passage SBK

To further assess the functional characteristics of SBK cultivated in Y-27632, we examined the global expression profile of early and late pass cells treated with IL-17 and IFN-γ, as this has been previously reported to stimulate an immunologic response in cultured keratinocytes [26]. Using a selection criteria of | log_2_ fold-change | > 2 and adjusted p value < 0.01, we identified 724 differentially expressed genes in the early passage group, and 1307 in the late passage (Fig 4A-B, S3 Appendix). Additionally, consistent with the previous report [26] we found human β-defensin 2 (βD2), 3 (βD3), and RNAase7 to be among the significantly upregulated genes in response to cytokine treatment in both the early and late passage groups. Next, we compared the overlap of the significantly differential genes from each group (Fig 4C), finding greater than 50% of the upregulated genes in early passage were represented in the late passage. However, this contrasted with less than 50% of the identified genes in late passage being similarly upregulated in early passage. We performed a chi-squared test to measure the overlap of genes between the two groups, yielding a p value <0.0001, indicating signficant concordance. This suggets that the molecular pathways of response to cytokine treatment is relatively preserved in late passage cells.

**Fig 4.**
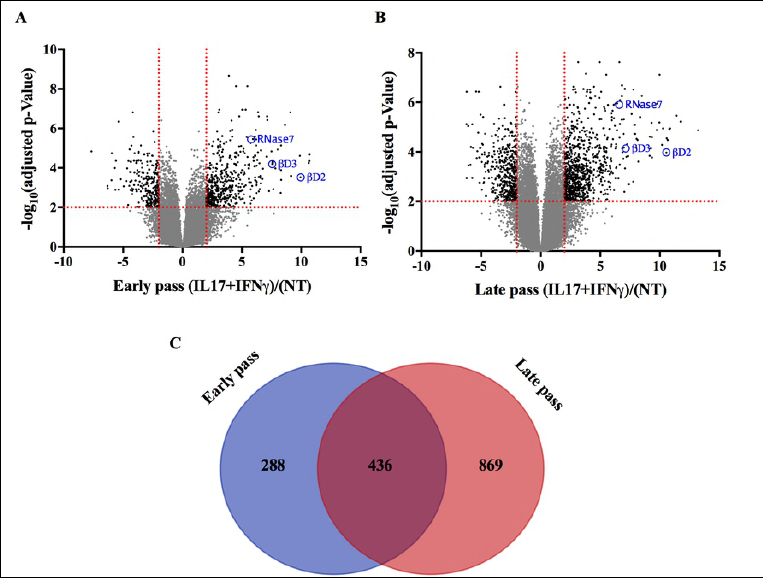
Functional characterization of early and late passage SBK. (A-B) RNA-seq analysis was was conducted on early and late passage SBK stimulated with IL-17 and IFN-γ for 48 hours. ANOVA analysis was performed, and the resulting -log_10_(adjusted p-value for false discovery) vs. log_2_ fold-change for difference of treatment was plotted. Values with | log_2_ fold-change | > 2 and an adjusted p value < 0.01 are highlighted in black (N=2 SBK patient cell lines, with 2 technical replicates for each treatment). (C) Venn diagram comparing the early and late passage significantly differential genes highlighted in Fig 4 A-B. Chi-square test was used to analyze contingency, resulting in a p <0.0001.

## Discussion

Improvement of *ex vivo* methods for culturing primary keratinocytes can allow further study of keratinocyte function in the setting of important cutaneous diseases such as methicillin-resistant *Staphylococcus aureus* abcesses, atopic dermatits, and psoriasis. Isolation techniques from neonatal foreskin keratinocytes are limited by sex exclusivity and temporal restrictions, surgical explants require a highly invasive procedure for collection, and skin biopsies confer small cellular yields. Further, all techniques require chemical separation of the epidermis from the dermis.

The suction blister method for isolating primary keratinoytes overcomes these limitations. However, the cells have proven difficult to culture due to their relatively quick differentiation and senecence *ex vivo* [4, 5]. Our results indicate that the addition of the Rho kinase (ROCK) inhibitor Y-27632 improves the proliferative capacity of the cells, increasing the population doubling from ∼2 to 15, thereby providing adequate cell numbers for experimentation.

Y-27632 does not fully immortalize the SBK, as evidenced by their senecence after ∼70 days in culture. Additionally, our results showing an increased IVL:KRT14 protein expression ratio in the late passage cells may allow this value to be used as a measure of the heterogeneity and functional senescence of the culture. If further validated, IVL:KRT14 or other protein expression ratios could aide researchers in standardizing cultures within and between experiments. The proliferative differences seen between our SBK and published reports with other epithelial cells grown indefinitely with Y-27632 [10, 27, 28] may be due to the variations intrinsic to different body sites. For example, foreskin-derived organotypic epithelia are more hyperplastic than trunk skin, and site-specific differences in keratin protein expression have been reported [29, 30]. Furthermore, the negative pressure of suction may impact epithelial-to-mesenchymal transition (EMT) pathways that influence differentiation [31]. Identifying and overcoming the mechanisms involved in the observed senescence would further improve the utility of this methodology to enable study of the role of keratinocytes in cutaneous disease and wound repair.

## Materials and Methods

### Patient Information

All human sample collection and processing were performed with approval of the National Institute of Allergy and Infectious Disease IRB, which approved the associated clinical trial (NCT02262819). All subjects gave written and verbal consent prior to sample collection. Participants ranged from 22-26 years of age, and comprised of 6 males and 1 female.

### Keratinocyte cell culture

Suction blister method keratinocytes were isolated as previously described [3]. Briefly, our eight-well suction chamber was placed onto the participants’ ethanol-sterilized volar forearm and placed under 30 mm Hg of suction for 2 hours using a micro-derm abrasion device (Kendal Diamond HB-SF02). After device removal, the resultant epidermal blister roofs were surgically removed and treated with trypsin to produce a single cell suspension. Culture expansion was performed as previously described [3, 10, 32] with irradiated 3T3-J2 fibroblasts in the presence or absence of 10μM Y-27632 (Tocris Bioscience). The human neonatal foreskin keratinocyte cell line was a generous gift from Dr. Alison McBride (NIH), and its isolation was described previously [10, 11]. The Feeder medium for the co-cultured cells contained 3:1 [v/v] Ham’s F-12 Nutrient mix (Invitrogen):DMEM high glucose, no glutamine (Invitrogen), 5% Newborn Calf Serum (Thermo Fisher Scientific), 0.4 μg/mL hydrocortisone (Sigma-Aldrich), 5 μg/mL insulin, 8.4 ng/mL Cholera Toxin, 10 ng/mL EGF (Invitrogen), 24 μg/mL adenine (Sigma-Aldrich), 10 U/mL Penicillin and 10 μg/mL streptomycin (Gibco), 2 mM L-glutamine (Gibco), 1X Primocin (Invitrogen). 3T3-J2 feeder fibroblasts were sub-cultured in DMEM (1X) – GlutaMAX^™^ (Invitrogen), 10% Fetal Bovine Serum (Hyclone), 25 mM HEPES (Corning), 50 μg/mL gentamicin (Quality Biological). The fibroblasts were cultured in 175cm^2^ flasks at an initial density of 2×10^6^ cells, and passed every 3 days using .25% trypsin-EDTA (Gibco) to detach cells. Prior to co-culture with the keratinocytes, 3T3-J2 cells were gamma-irradiated with a dose of 6000 Rads using a Co-60 source. Co-cultured cells were passed using Versene first to detach fibroblast feeder cells, followed by .25% trypsin-EDTA to detach keratinocytes. After detachment, keratinocytes were passed through a 70 μM nylon filter to break up cell aggregates, and then seeded into a 10 cm dish at a density of 4×10^5^ keratinocytes:1×10^6^ 3T3-J2 fibroblasts. Population Doubling (PD) was calculated using the equation PD = 3.32(log[number of cells harvested/number of cells seeded]).

### Western blot analysis

To measure the proportion of differentiated keratinocytes in culture, early passage cells (P4 and P5) were detached as described above, counted, and added to 12 well plates at a density of 1×10^5^/well, on top of 2×10^5^ irradiated 3T3-J2 fibroblasts. The media used was either complete F-medium, or F-medium lacking Y-27632. At Day 0, 4, 8, 12, and 16 cells were washed with PBS, followed by Versene, followed again by 2 PBS washes. Protein was then extracted using RIPA buffer (Sigma) according to the manufacturer’s procedure and quantified using the Pierce BCA assay. 17 ng of protein was loaded onto a 4-12% Bis-tris gel (Gibco), transferred onto a nitrocellulose membrane and stained with the following primary antibodies: mouse monoclonal anti-Involucrin (SY5) (Thermo Fisher, MA511803), 1:200 dilution; goat polyclonal anti-Cytokeratin 14 (LL001) 1:200 dilution (Santa Cruz Biotechnology, sc-53253); rabbit polyclonal anti-GAPDH, 1:500 dilution (Cell Signaling, 88845). The secondaries used were anti-goat 680 conjugated (Rockland, 605-730-002), 1:10,000 dilution, anti-mouse 800 conjugated (Rockland, 610-132-003), 1:10,000 dilution; anti-rabbit 800 conjugated (Rockland, 611-145-003) 1:10,000 dilution. The 9120 Odyssey (LI-COR) system was used to image the blots. Two independent experiments were performed, with n=2 cell lines tested total. Integrated densitometry was performed with ImageJ (NIH), and the ratio of IVL/KRT14 was calculated. GAPDH was included as a loading control.

The IVL/KRT14 ratio was also measured for early and late passage SBK cultures. Early (passage 4-5) and late (passage 12-14) cells were seeded in 12 well culture plates with the above described feeder system at a density of 1×10^5^ keratinocytes and 4×10^5^ irradiated fibroblasts. After 3 days the media was aspirated and the cells were washed 3X and treated with Versene to remove the fibroblast feeder layer. Protein was isolated and quantified as described in the previous paragraph. N=4 patient cell lines were tested, and a parametric paired t test was used to measure significance, with P<0.05 to measure significance.

### Cell size quantification

Keratinocytes were imaged with a Zeiss Axiovert 200 using an exposure time of 700 ms. The magnification used to image the cells was 10X, and the cell size was quantified using ImageJ. n=2 patients were examined, and 3 randomly selected fields of each culture were used to quantify the size, measuring 25 cells/field. HFK were measured similarly except that only one cell line was used for quantification.

### RNA-seq analysis

To characterize the effect of Y-27632 on the expression profile of SBK, we incubated cells starting from isolation in the presence (+Y) or absence (-Y) of the inhibitor in the co-culture feeder system described early in the methods. After removal of the feeder cells with Versene, RNA from SBK at passage 4 and 5 for the +Y group and 4, 5, and 6 or the -Y group was isolated using the RNeasy kit (Qiagen).

To compare the differential transcriptional response to cytokine treatment between early and late keratinocytes passaged with Y-27632 we adapted the IL-17 + IFN-γ stimulation conditions described previously [26] with some modifications. Two types of media were used: SFM (+), which contains SFM medium (Gibco) with 10 ng/mL EGF (Gibco), 50 μg/mL Bovine Pituitary Extract (Gibco), 1X primocin, and 10 μM Y-27632; or SFM(-), which only has SFM medium with 50 μg/mL Bovine Pituitary Extract and 1X primocin. We took early (pass 4) and late (pass 12) keratinocytes out of culture and seeded them into a 24 well cell culture plate with 1 mL of SFM(+) at a density of 2×10^5^ cells/well and incubated 37° C and 5% CO_2_ for 24 hours. The next day, cells were washed 3X with PBS and 1 mL of SFM(-) was added. Cells were incubated at 37°C and 5% CO_2_ for 24 hours. The next day, cells were washed with PBS, and 400 μL of SFM(-) was added containing 10 ng/mL of IL-17A (R&D Systems), IFN-γ (R&D Systems), or 0.1% BSA in PBS for the no treatment control. Cells were incubated at 37°C and 5% CO_2_ for 48 hours, after which RNA was isolated using the RNeasy kit (Qiagen). N=2 early and late pass patient cell lines were tested, with two technical replicates for each condition.

RNA was assessed for quality and degradation on the Agilent 2200 Tapestation System (Agilent, Santa Clara, CA), using the product RNA ScreenTape (Catalog # 5067-5576). All samples scored RINe values ≥9.5. RNA purity and concentration was assessed using the Nanodrop One UV-Vis Spectrophotometer (ThermoFischer Scientific, Waltham, MA).

Library preparation was completed using the Illumina Neoprep Library Preparation System (Illumina, San Diego, CA). The product TruSeq Stranded mRNA Library Prep Kit for NeoPrep (Catalog # NP-202-1001) was used for library construction using 100 ng Total RNA input per sample, following Illumina protocol “TruSeq Stranded mRNA Library Prep for NeoPrep Reference Guide” (Document # 15049725 v03, June 2016). TruSeq Stranded mRNA Protocol Version 1.1.7.6 was used, with default settings of 200 bp target insert size and 15 cycles PCR.

Library validation was performed on the Agilent 2200 Tapestation System (Agilent, Santa Clara, CA), using the product High Sensitivity D1000 ScreenTape (Catalog # 5067-5584) to verify library size and purity. Library quantification via qPCR was performed on the Applied Biosystems 7900HT Fast Real-Time PCR System (ThermoFischer Scientific, Waltham, MA) using KAPA Biosystems Complete Library Quantification Kit for Illumina (Catalog # KK4835).

Libraries were normalized to 2 nM concentration. Equimolar aliquots of 5ul of each library were then pooled. The 2nm library pool was then denatured and diluted to 1.8 pM, following the Illumina protocol “NextSeq System Denature and Dilute Libraries Guide” (Document # 15048776 v02, January 2016). PhiX was added at 1% to serve as an internal control. The resultant final library pool was 1.8 pM final concentration with 1% PhiX spike-in.

Single-end sequencing was completed on an Illumina NextSeq 500 system, running Illumina NextSeq Control Software System Suite version 2.1.2 and RTA version 2.4.11. The final library pool was sequenced via 1 × 76 bp run configuration using the product NextSeq 500/550 High Output v2 kit,75 cycles (Catalog # FC-404-2005).

Sequencing reads (single-end, 75 basepair) where aligned to human genome (version hg38 downloaded from Ensemble) using Qiagen CLC Genomics Workbench version 11.0.1 in strand-specific mode with a maxiumum of 10 hits per read. Non-specific matches were assigned via the EM algorithm. Total_exon_reads where taken as the measure of raw counts per locus.

Raw counts inflated by 1 and transformed by log2 before upper quartile scaling normalization and differential expression testing using JMP/Genomics (SAS Institute, Cary NC) version mixed effects ANOVA.

### Live imaging time lapse video

SBK were isolated from one patient and seeded in a 24 well cell culture plate in the co-culture system as described above. The plate was incubated in an insulated container on a movable stage that was kept at 37°C and 5% CO_2_, and the cells were imaged with a Leica Inverted Epifluorescent Microscope equipped with Leica DFC 345 monochrome camera for Bright Field and DIC images. The camera position for each well was programmed using LAS X, and a 10X picture was taken every 10 minutes for 43 hours. The images were compiled into a movie using Imaris 9.0.

### Organotypic raft cultures

The procedure was performed as previously described [33] and modified for our purposes. 2.0×10^5^ J2-3T3 fibroblasts were seeded in 100% type 1A rat-tail collagen (Corning) with 2×10^5^ keratinocytes of indicated passage number in a 24 well cell culture plate with 1mL of raft medium [3:1 Ham’s F12 nutrient mix (Invitrogen): DMEM high glucose no glutamine (Invitrogen), 5% Newborn calf serum (Thermo Fisher Scientific), 0.4 μg/mL hydrocortisone (Sigma-Aldrich), 10 ng/mL human EGF (Invitrogen), 0.1 nM Cholera Toxin, 5 μg/mL insulin, 2mM glutamine (Gibco), 1X Primocin (Invitrogen)] and incubated over night at 37°C and 5% CO_2_. The next day, rafts were lifted onto stainless steel metal mesh and incubated in 6 well cell plates at the liquid air interface for 13 days. ∼3mL of raft medium was used for each raft and changed every 3 days. After 13 days, the rafts were washed with PBS and fixed in formalin and processed as described below.

### Histopathology

Organotypic skin culture were fixed in 10% neutral buffered formalin, sectioned, and blocked in paraffin for histological analysis. Tissue sections (5 μm) were stained with hematoxylin and eosin (H&E) for routine histopathology and then evaluated by a pathologist in a blinded manner. Sections were examined under light microscopy using an Olympus BX51 microscope and photographs were taken using an Olympus DP73 camera.

### Immunohistochemistry (IHC) & Immunofluorescence (IF)

For immunohistochemical (IHC) and immunofluorescence (IF) analyses, tissue sections (5 μm) were heated to 60°C for 1 hour, deparaffinized with xylene washes, and rehydrated with alcohol-graduated washes. Heat-induced epitope retrieval (HIER) was performed using Epitope Retrieval Solution 1, pH 6.0, heated to 100°C for 20 minutes. The slides were then incubated with hydrogen peroxide to quench endogenous peroxidase activity prior to applying the primary antibody. For IHC, detection was performed with DAB chromogen and counterstaining with hematoxylin using the Bond RX (Leica Biosystems). For immunofluorescence (IF), tissue slides were first treated with a mouse monoclonal Involucrin (SY5) (Thermo Fisher Scientific, MA511803) primary antibody at a dilution of 1:80, labeled with an Alexa Fluor 594 conjugated goat anti-mouse (Thermo Fisher Scientific, A-11032) secondary antibody. This was followed by application of a second primary antibody, a goat polyclonal Cytokeratin 14 (C-14) (Santa Cruz Biotechnology, sc-53253) primary antibody at a dilution of 1:50 and labeled with an Alexa Fluor 488 conjugated donkey anti-goat (Thermo Fisher Scientific, A-11055) secondary antibody. Both secondary fluorescent antibodies were used at a dilution of 1:500. Nuclei were counterstained with DAPI (Vector Laboratories) and sections were mounted with ProLong Gold anti-fade reagent (Invitrogen). Sections were examined by light microscopy using an Olympus BX51 microscope and photomicrographs were taken using an Olympus DP73 camera.

## Statistical analysis

GraphPad Prism 7 (Graphpad Software Inc., La Jolla CA) was used for non-RNAseq statistical analysis. To measure the significant differences between cytoplasmic areas we used a one-way ANOVA with Sidak’s test for multiple comparisons; an adjusted p value of <0.05 was considered significant. To measure significant differences between early and late pass IVL/KRT14 ratios we used a parametric paired t test.

## Author contributions

EDA, IS, SKD, and IAM designed the studies. EDA, IS, NE performed experiments and analyzed data. MM and INM contributed the IHC and IF figures, and FOC and TGM ran and analyzed the RNAseq data. CAM designed the 3D suction blister device. EDA wrote the manuscript, and IAM, NE, IS, and SKD helped edit. All authors critically reviewed the manuscript.

## Acknowledgements

This research was supported by the Intramural Research Program of the NIH and the National Institute of Allergy and Infectious Diseases. We thank Alison McBride for guidance on establishing keratinocyte culture techniques and gifting us the HFK cell line used in this study. We also thank Sundar Ganesan, Hatice Karauzum, Arhum Saleem, Mark Kieh, Wei Chen, Aaron Robertson, Pamela Welch, and Dirk Darnell for technical and administrative assistance.

## Supporting information

**S1 Fig. Y-27632 increases the proliferative capacity of HFK.** Sum total of HFK generated from the +Y (black) and -Y (red) conditions after 70 and 25 days respectively.

**S2 Fig. Y-27632 causes SBK to clump and form colonies more rapidaly immediately after isolation.** Time lapse of SBK cultured 1.5 hours after isolation from the patient, and with (+Y) or without (-Y) the Y-27632 inhibitor and monitored for colony formation. The dotted circles highlight colonies that are beginning to form after 43 hours in the +Y condition.

**S3 Fig. Y-27632 delays the expression of late stage differentiation proteins, but late pass SBK present a heterologous stratification pattern.** (A) KRT14 and IVL expression levels in SBK seeded in culture dishes and incubated with or without Y-27632 for 0, 4, 8, 12, or 16 days. The ratio of IVL/KRT14 ratios were calculated by integrated densitometry with ImageJ and normalized to GAPDH expression (right, n=2 SBK patient cell lines were tested at passage 5). (B) KRT14 and IVL expression levels in early and late passage SBK cultured with Y-27632. Normalized IVL/KRT14 ratios are presented (right, n=4,and each shape represents SBK from a different patient). ^*^, p<0.05

**S1 Appendix. Data file for 3D Printing Suction Blister Device.** (To be uploaded upon manuscript acceptance).

**S2 Appendix. RNA-seq Differential genes +Y27632 vs -Y27632.** The differential response to Y-27632 treatment was determined and genes with | log_2_ fold-change | > 2.6 and an adjusted p value <0.5 are listed in the first tab of this document. The second tab presents the lists genes used to construct the venn diagram in Fig 2B. Full gene list will be deposited onto Geo database (accession number TBD).

**S3 Appendix. RNA-seq Differential genes Late pass (IL17+IFNG/NT) vs Early pass (IL17+IFNG/NT).** The differential response to cytokine treatment (IL17+IFNγ vs. NT) was determined independently for early and late pass cultures, and all genes with an adjusted p value <0.01 and | log_2_ fold-change | > 2 for each group are listed in this document. Full gene list will be deposited onto Geo database (accession number TBD).

**S1 Movie. SBK time lapse video 1.5 hours after isolation.** SBK were seeded 1.5 hours after isolation from blister roofs and incubated either with or without Y-27632 in the co-culture feeder system for 43 hours.

